# Experimental Validation of an Organ-on-Chip for Mechanical Stimulation of Cell Cultures

**DOI:** 10.1101/2023.12.15.571835

**Authors:** Maria Testa, Salvatore Tornabene, Sofia Di Leonardo, Gaetano Burriesci, Vincenzo La Carrubba, Francesco Lopresti

## Abstract

This paper focuses on the design, fabrication, and characterization of a platform aiming to dynamically culture cells under mechanical stimulation. The platform, made of polymethylmethacrylate (PMMA), allows real-time mechanical stimuli, providing valuable insights for living tissue models. Through mechanical testing and dynamic microfluidic tests, the chip functionality was assessed. The experimental results validation showcases the potential of the device in mimicking physiological conditions, offering a promising avenue for pharmaceutical testing advancement.

## Introduction

Pharmaceutical research strives to enhance human health through the identification and development of new medicinal compounds and therapies. This intricate process involves exploring fundamental aspects to identify potential therapeutic targets. The subsequent pre-clinical phase, which includes *in vitro* and animal model testing, aims to assess the efficacy, safety, and toxicity of compounds. However, transitioning from pre-clinical to clinical phases poses challenges due to discrepancies in results, primarily attributed to the limitations of *in vitro* models in accurately reflecting complex human physiology. Amidst growing societal and regulatory pressure to reduce animal testing, innovative technologies such as Organ-on-Chips (OoCs) have emerged [1]–[3].

Advanced OoC systems can also deliver mechanical, chemical, electrical, and flow stimuli on the cultured cells. OoCs provide controlled mechanical stimuli, including shear force, traction and compression, influencing the cells within organs. Chemical stimuli involve flowing substances through microfluidic channels to mimic *in vivo* conditions. Controlled electrical stimuli allow the study of electrostimulation effects on organs and their self-regulatory behavior. Microfluidic flows deliver nutrients and chemicals, replicating physiological conditions and applying mechanical stimulation, such as shear stress [4]–[21].

In this work, a novel design of an OoC microfluidic platform which combines mechanical and shear stress stimuli to create multi-stimulation states is presented. This device is capable of imposing cyclic deformations and stresses through a deformable diaphragm embedded in a rigid structure. The final application involves placing a flexible cellularized scaffold on top of the diaphragm to administer physiological mechanical stimuli to cell cultures.

## Material and method

The following materials were used in the fabrication of this device: PMMA (Clarex, nitto Jushi Kogyo Co. Ltd, JP), Pure Ethanol (Et-OH) (Purchased from Sigma-Aldrich) and Silicone (Ultra-thin - High precision, 100 µm thick purchased from MyTech Ltd., Germany).

The PMoC was designed to measure the volume of fluid displaced by the diaphragm’s deformation under inlet pressure. PMMA, chosen for its thermoplastic and amorphous properties, served as the chip material. This transparent and readily available polymer was processed using rapid prototyping techniques such as laser cutting and multilayer assembling [9], [18], [19].

The device features separate chambers divided by a diaphragm of silicone that deforms under externally generated pressure, providing mechanical stimulation to cultured cells. The fluid-containing chamber is connected to a calibrated capillary for displaced volume measurement. The design includes inlets for fluid and pressure application, as well as outlets for bubble elimination and pressure adjustment. Layers were fabricated according to design using AutoLaser software for a CO_2_ Laser Cutter (Maitech, 40 W). After parameter optimization tests, layers were processed accordingly to the literature [23]–[26].

Assembling the layers involved a hydraulic press with heated plates, ensuring proper alignment with alignment pins in a specially made aluminum mold. Prior to assembly, ethanol was deposited to cover contact surfaces, acting as a weak solvent for PMMA and facilitating efficient assembly while preserving the integrity of drilled microchannels. The layers are put in order and stacked inside the aluminum mold for hydraulic press (Figure1).

**Figure 1:**
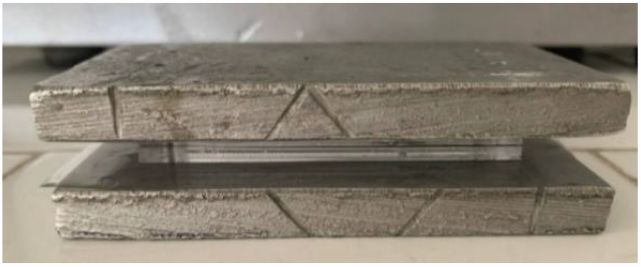
Chip stacked inside the aluminum mold

Assembly parameters, such as temperature (70 °C) and pressure (8000 kPa), were chosen based on literature recommendations and maintained for 3 minutes to achieve a successful fabrication process [24]–[26].

To determine the mechanical properties of the silicone, uniaxial and biaxial mechanical tests were conducted. Uniaxial characterizations were performed on 6 specimens for each sample by means of an Instron 5943 Series single-column universal mechanical characterization apparatus, while biaxial characteristics were assessed with a Cell Scale Biotester 5000. Uniaxial test parameters were obtained following BS ISO 37:2011, utilizing type 3 dog-bone specimens with a 16 mm gauge length and width of 4 mm, tested in displacement control at 200 mm/min [27]– [30]. For biaxial analysis, 15 specimens of gauge area of 10×10 mm^2^ were tested using a BioRakes-anchored setup in displacement control at a speed of 3.3 mm/min. Black matte varnish facilitated Digital Image Correlation (DIC) analysis for strain mapping. The DIC analysis focused on the ramp-up phase at 30% strain for all 15 specimens. Images of deformed specimens were acquired every second, generating strain maps by using the MATLAB open-source code Ncorr [31].

To evaluate pressures and flows during PMoC operation, pressure and flow sensors were employed. Microfluidic pressure sensors (LabSmith, uPS-series) measured pressures, affixed above and below the diaphragm. A Sensirion SLF3S-0600 flow sensor, coupled with a programmable syringe pump (Lead Fluid TYD01), measured flows. The syringe pump delivered colored liquids into the device, allowing visualization of deviations within the capillary and creating a pressure difference affecting the membrane.

## Results and Discussion

Figure 2 shows the device after the rapid prototyping fabrication, ready for experimental test.

**Figure 2:**
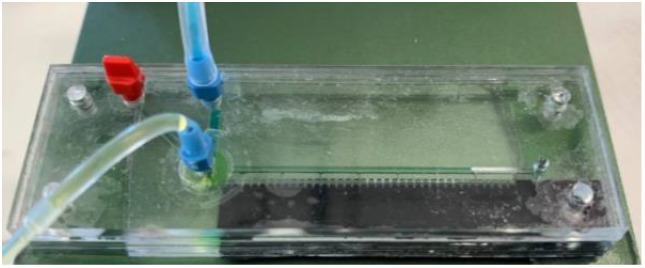
Left top view of the device

The mechanical characterization of the silicone film (100 µm thick) with uniaxial tensile tests gave the following results:

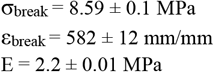

Following the analysis of data from uniaxial silicone film emerged as an adequate choice for the diaphragm within the chip. Consequently, biaxial mechanical testing was conducted on the same material. The acquired data were subsequently employed in a Digital Image Correlation (DIC) analysis. This process allowed the generation of a sequence of strain maps (Figure 3).

**Figure 3:**
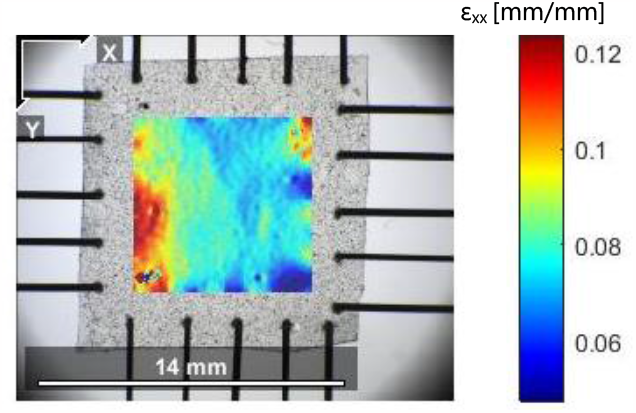
Deformation Map provide by N_corr_

The DIC analysis process unfolds in multiple stages. Firstly, a reference image is established, and subsequent photos are loaded. In this instance, the eighth photo was selected as the initial instant, extending through the subsequent images up to the peak strain, with an applied pre-load of 0.5 N. Choosing for a photo not precisely corresponding to the initial one is akin to applying a preload, a deliberate choice reflecting the preloaded state of the aperture between the chip layers. Subsequently, an appropriate Region of Interest (ROI) of 90% gauge area and DIC parameters are defined for further analysis. The material is isotropic so the stress and strain are mediated in the two directions.

The result shows a slightly higher elastic modulus (evaluated at low strains) than that determined with the uniaxial test, *E*_*uni*_ *=* 2.2 *MPa E*_*biax*_ *=* 4.3 *MPa (*Figure 4). This result was expected and has already been reported in the literature for materials with similar behavior [27]–[35].

**Figure 4:**
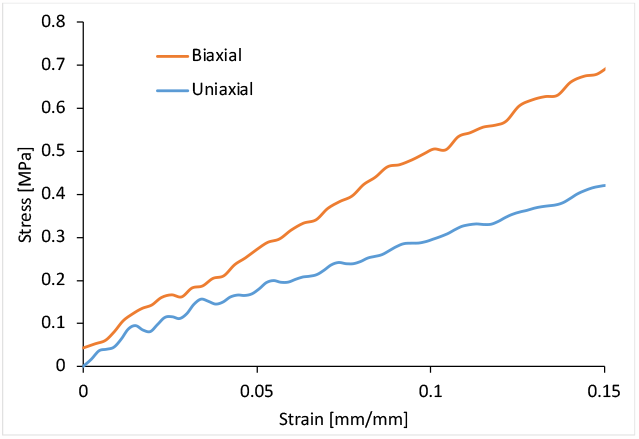
Plot average stress-strain curves of silicone deformation

Pressure and flow sensor data in the main chamber responsible for diaphragm deformation were sampled at 100 µs intervals at approximately 28 °C. The cyclic stimulation by triangular shape, with a 60-second period, displays a regular pattern with linear growth up to a peak of 28 kPa in the loading phase, followed by a linear decrease to 5 kPa in the unloading phase (Figure 5). The subsequent pressure trend shows slight deviations from linearity between 5 kPa and 0 kPa. The flow rate trend, though more irregular, exhibits an observable peak of approximately 120 µL/min compared to the set 200 µL/min, attributed to coupling issues in the delivery system. Despite this, the device manages to return to zero.

**Figure 5:**
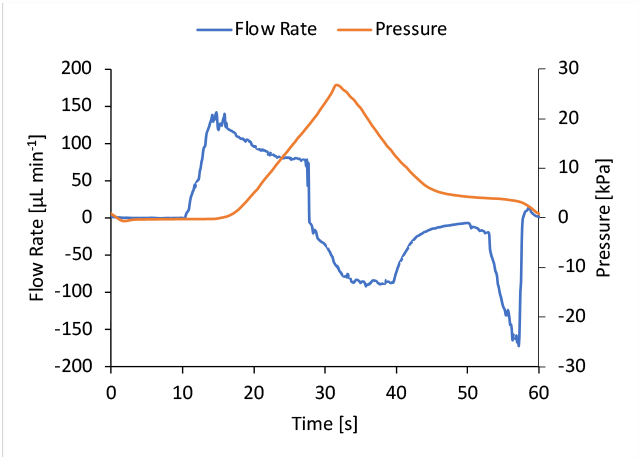
Pressure and flow graph recorded for the device

Additionally, measurements of pressure in the lower chamber confirmed that stimulation arises solely from the flow rate change in the upper chamber, with a peak of pressure of about -0.12 kPa, significantly lower than the upper chamber. The displacement of the liquid head in the capillary was analyzed for quantitative evaluation of the volume displaced during diaphragm delivery of the inlet pressure. The maximum displacement caused by diaphragm deformation (ΔL_max_) is 31 mm (Figure 6), which corresponds to a volume equal to 15.05 mm^3^, thus giving an experimental information about the moved volume by the diaphragm deformation.

**Figure 6:**
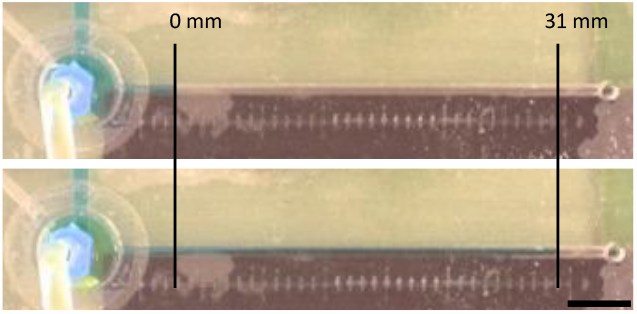
Picture representing the movement of the fluid in the capillary of the chip. Scale bar is 5 mm

## Conclusion

The presented paper shows the design and fabrication of an Organ-on-Chip (OoC) device incorporating a deformable diaphragm. Results indicate the successful creation of an OoC system by assembling PMMA layers of varying thicknesses, introducing a deformable system for delivering mechanical stimuli to cells. Experimental analyses demonstrated the effectiveness of this mechanical stimulus delivery.

In summary, the paper introduces an affordable, easily reproducible, and user-friendly OoC system adaptable to diverse cell cultures by controlling membrane deformation. However, further development is necessary, including coupling the deformable diaphragm with a suitable scaffold for cell cultures and conducting dynamic tests on mechanical stimulus delivery.

The evidence derived from this work suggests broad applications for the OoC system, enabling testing of various cell types on a single off-the-shelf platform by adjusting diaphragm pressures and frequencies. The study demonstrates the attainment of high levels of efficiently regulated mechanical stimulation through input flow rate adjustments to the device. Future perspectives involve enhancing the delivery system to achieve greater control over stimulations delivered to diverse cell types and a numerical analysis for a computational validation.

## Bibliography

[1] J. A. DiMasi, H. G. Grabowski, and R. W. Hansen, “Innovation in the pharmaceutical industry: New estimates of R&D costs,” J Health Econ, vol. 47, pp. 20–33, May 2016, doi: 10.1016/j.jhealeco.2016.01.012.

[2] C. M. Leung et al., “A guide to the organ-on-a-chip,” Nature Reviews Methods Primers, vol. 2, no. 1. Springer Nature, Dec. 01, 2022. doi: 10.1038/s43586-022-00118-6.

[3] L. A. Low, C. Mummery, B. R. Berridge, C. P. Austin, and D. A. Tagle, “Organs-on-chips: into the next decade,” Nature Reviews Drug Discovery, vol. 20, no. 5. Nature Research, pp. 345–361, May 01, 2021. doi: 10.1038/s41573-020-0079-3.

[4] A. Mainardi, P. Occhetta, I. Martin, M. Rasponi, and A. Barbero, “Towards Modelling The Joint On A Chip: A Mechanically Active Microfluidic System Designed For Engineered 3d Multi-Layer Osteochondral Tissues,” Osteoarthritis Cartilage, vol. 31, pp. S56–S57, Mar. 2023, doi: 10.1016/j.joca.2023.01.551.

[5] E. Ferrari, C. Palma, S. Vesentini, P. Occhetta, and M. Rasponi, “Integrating Biosensors in Organs-on-Chip Devices: A Perspective on Current Strategies to Monitor Microphysiological Systems,” Biosensors (Basel), vol. 10, no. 9, p. 110, Aug. 2020, doi: 10.3390/bios10090110.

[6] E. Ferrari and M. Rasponi, “Liver–Heart on chip models for drug safety,” APL Bioeng, vol. 5, no. 3, Sep. 2021, doi: 10.1063/5.0048986.

[7] S. Schneider, D. Gruner, A. Richter, and P. Loskill, “Membrane integration into PDMS-free microfluidic platforms for organ-on-chip and analytical chemistry applications,” Lab on a Chip, vol. 21, no. 10. Royal Society of Chemistry, pp. 1866–1885, May 21, 2021. doi: 10.1039/d1lc00188d.

[8] P. I. C. Claro, A. R. S. Neto, A. C. C. Bibbo, L. H. C. Mattoso, M. S. R. Bastos, and J. M. Marconcini, “Biodegradable Blends with Potential Use in Packaging: A Comparison of PLA/Chitosan and PLA/Cellulose Acetate Films,” J Polym Environ, vol. 24, no. 4, pp. 363–371, Dec. 2016, doi: 10.1007/s10924-016-0785-4.

[9] J. H. Huang et al., “A microfluidic method to measure bulging heights for bulge testing of polydimethylsiloxane (PDMS) and polyurethane (PU) elastomeric membranes,” RSC Adv, vol. 8, no. 38, pp. 21133–21138, 2018, doi: 10.1039/c8ra01256c.

[10] C. R. Wan, S. Chung, and R. D. Kamm, “Differentiation of embryonic stem cells into cardiomyocytes in a compliant microfluidic system,” Ann Biomed Eng, vol. 39, no. 6, pp. 1840–1847, Jun. 2011, doi: 10.1007/s10439-011-0275-8.

[11] D. Bavli et al., “Real-time monitoring of metabolic function in liver-onchip microdevices tracks the dynamics of Mitochondrial dysfunction,” Proc Natl Acad Sci U S A, vol. 113, no. 16, pp. E2231–E2240, Apr. 2016, doi: 10.1073/pnas.1522556113.

[12] W. E. Sinclair et al., “Development of microfluidic platform that enables ‘on-chip’ imaging of cells exposed to shear stress and cyclic stretch,” Microfluid Nanofluidics, vol. 27, no. 2, Feb. 2023, doi: 10.1007/s10404-022-02619-y.

[13] D. Huh et al., “Acoustically detectable cellular-level lung injury induced by fluid mechanical stresses in microfluidic airway systems,” 2007. [Online]. Available: https://www.pnas.orgcgidoi10.1073pnas.0610868104

[14] J. D. Stucki et al., “Medium throughput breathing human primary cell alveolus-on-chip model,” Sci Rep, vol. 8, no. 1, Dec. 2018, doi: 10.1038/s41598-018-32523-x.

[15] A. Bein et al., “Microfluidic Organ-on-a-Chip Models of Human Intestine,” CMGH, vol. 5, no. 4. Elsevier Inc, pp. 659–668, Jan. 01, 2018. doi: 10.1016/j.jcmgh.2017.12.010.

[16] K. Schimek et al., “Integrating biological vasculature into a multi-organ-chip microsystem,” Lab Chip, vol. 13, no. 18, pp. 3588–3598, Sep. 2013, doi: 10.1039/c3lc50217a.

[17] D. Bavli et al., “Real-time monitoring of metabolic function in liver-onchip microdevices tracks the dynamics of Mitochondrial dysfunction,” Proc Natl Acad Sci U S A, vol. 113, no. 16, pp. E2231–E2240, Apr. 2016, doi: 10.1073/pnas.1522556113.

[18] W. J. Polacheck, R. Li, S. G. M. Uzel, and R. D. Kamm, “Microfluidic platforms for mechanobiology,” Lab on a Chip, vol. 13, no. 12. Royal Society of Chemistry, pp. 2252–2267, Jun. 21, 2013. doi: 10.1039/c3lc41393d.

[19] K. Kaarj and J. Y. Yoon, “Methods of delivering mechanical stimuli to Organ-on-a-Chip,” Micromachines, vol. 10, no. 10. MDPI AG, Oct. 01, 2019. doi: 10.3390/mi10100700.

[20] O. T. Guenat and F. Berthiaume, “Incorporating mechanical strain in organs-on-a-chip: Lung and skin,” Biomicrofluidics, vol. 12, no. 4, Jul. 2018, doi: 10.1063/1.5024895.

[21] J. A. Terrell, C. G. Jones, G. K. M. Kabandana, and C. Chen, “From cells-on-a-chip to organs-on-a-chip: scaffolding materials for 3D cell culture in microfluidics,” J Mater Chem B, vol. 8, no. 31, pp. 6667–6685, 2020, doi: 10.1039/D0TB00718H.

[22] J. M. de Hoyos-Vega, A. M. Gonzalez-Suarez, and J. L. Garcia-Cordero, “A versatile microfluidic device for multiple ex vivo/in vitro tissue assays unrestrained from tissue topography,” Microsyst Nanoeng, vol. 6, no. 1, Dec. 2020, doi: 10.1038/s41378-020-0156-0.

[23] A. E. Ongaro et al., “Laser Ablation of Poly(lactic acid) Sheets for the Rapid Prototyping of Sustainable, Single-Use, Disposable Medical Microcomponents,” ACS Sustain Chem Eng, vol. 6, no. 4, pp. 4899–4908, Apr. 2018, doi: 10.1021/acssuschemeng.7b04348.

[24] F. Lopresti, I. Keraite, A. E. Ongaro, N. M. Howarth, V. La Carrubba, and M. Kersaudy-Kerhoas, “Engineered Membranes for Residual Cell Trapping on Microfluidic Blood Plasma Separation Systems: A Comparison between Porous and Nanofibrous Membranes,” Membranes (Basel), vol. 11, no. 9, p. 680, Aug. 2021, doi: 10.3390/membranes11090680.

[25] F. Lopresti et al., “Core-shell PLA/Kef hybrid scaffolds for skin tissue engineering applications prepared by direct kefiran coating on PLA electrospun fibers optimized via air-plasma treatment,” Materials Science and Engineering: C, vol. 127, p. 112248, Aug. 2021, doi: 10.1016/j.msec.2021.112248.

[26] F. Lopresti, I. Keraite, A. E. Ongaro, N. M. Howarth, V. La Carrubba, and M. Kersaudy-Kerhoas, “Engineered Membranes for Residual Cell Trapping on Microfluidic Blood Plasma Separation Systems: A Comparison between Porous and Nanofibrous Membranes,” Membranes (Basel), vol. 11, no. 9, p. 680, Aug. 2021, doi: 10.3390/membranes11090680.

[27] A. Avanzini and D. Battini, “Integrated Experimental and Numerical Comparison of Different Approaches for Planar Biaxial Testing of a Hyperelastic Material,” Advances in Materials Science and Engineering, vol. 2016, pp. 1–12, 2016, doi: 10.1155/2016/6014129.

[28] A. Eilaghi, J. G. Flanagan, G. W. Brodland, and C. R. Ethier, “Strain Uniformity in Biaxial Specimens is Highly Sensitive to Attachment Details,” J Biomech Eng, vol. 131, no. 9, Sep. 2009, doi: 10.1115/1.3148467.

[29] H. Fehervary, M. Smoljkić, J. Vander Sloten, and N. Famaey, “Planar biaxial testing of soft biological tissue using rakes: A critical analysis of protocol and fitting process,” J Mech Behav Biomed Mater, vol. 61, pp. 135–151, Aug. 2016, doi: 10.1016/j.jmbbm.2016.01.011.

[30] G. M. Cooney, K. M. Moerman, M. Takaza, D. C. Winter, and C. K. Simms, “Uniaxial and biaxial mechanical properties of porcine linea alba,” J Mech Behav Biomed Mater, vol. 41, pp. 68–82, Jan. 2015, doi: 10.1016/j.jmbbm.2014.09.026.

[31] J. Blaber, B. Adair, and A. Antoniou, “Ncorr: Open-Source 2D Digital Image Correlation Matlab Software,” Exp Mech, vol. 55, no. 6, pp. 1105–1122, Jul. 2015, doi: 10.1007/s11340-015-0009-1.

[32] N. N. Azmi, M. N. A. Ab Patar, S. N. A. Mohd Noor, and J. Mahmud, “Testing standards assessment for silicone rubber,” in 2014 International Symposium on Technology Management and Emerging Technologies, IEEE, May 2014, pp. 332–336. doi: 10.1109/ISTMET.2014.6936529.

[33] S. N. A. M. Noor and J. Mahmud, “Modelling and Computation of Silicone Rubber Deformation Adapting Neo-Hookean Constitutive Equation,” in 2015 Fifth International Conference on Communication Systems and Network Technologies, IEEE, Apr. 2015, pp. 1323–1326. doi: 10.1109/CSNT.2015.276.

[34] A. Avanzini and D. Battini, “Integrated experimental and numerical comparison of different approaches for planar biaxial testing of a hyperelastic material,” Advances in Materials Science and Engineering, vol. 2016, 2016, doi: 10.1155/2016/6014129.

[35] S. Di Leonardo, A. Monteleone, P. Caruso, H. Meecham-Garcia, G. Pitarresi, and G. Burriesci, “Effect of the apron in the mechanical characterisation of hyperelastic materials by means of biaxial testing: A new method to improve accuracy,” J Mech Behav Biomed Mater, p. 106291, Dec. 2023, doi: 10.1016/j.jmbbm.2023.106291.

